# *C. elegans* LIN-28 controls temporal cell-fate progression by regulating LIN-46 expression via the 5’UTR of *lin-46* mRNA

**DOI:** 10.1101/697490

**Authors:** Orkan Ilbay, Charles Nelson, Victor Ambros

## Abstract

Human Lin28 is a conserved RNA-binding protein that promotes proliferation and pluripotency and can be oncogenic. Lin28 and *C. elegans* LIN-28 bind to precursor RNAs of the conserved, cellular differentiation-promoting, microRNA *let-7* and inhibits production of mature *let-7* microRNA. Lin28/LIN-28 also binds to and regulates many mRNAs in various cell types. However, the determinants and consequences of these LIN-28-mRNA interactions are not well understood. Here, we report that LIN-28 in *C. elegans* represses the expression of LIN-46, a downstream protein in the heterochronic pathway, via the 5’ UTR of the *lin-46* mRNA. We show that both LIN-28 and the 5’UTR of *lin-46* are required to prevent LIN-46 expression in the L1 and L2 stages, and that precocious LIN-46 expression is sufficient to skip L2 stage proliferative cell-fates, resulting in heterochronic defects similar to the ones observed in *lin-28(0)* animals. We propose that the *lin-46* 5’UTR mediates LIN-28 binding to and repression of the *lin-46* mRNA. Our results demonstrate that precocious LIN-46 expression alone can account for *lin-28(0)* phenotypes, demonstrating the biological importance of regulation of individual target mRNAs by LIN-28.

## INTRODUCTION

Animal development involves complex cell lineages within which different cell-fates are executed in specific orders and at a pace that is in synchrony with overall developmental rate. Through the regulated expression of, on the one hand, symmetric cell-fates that allow cell proliferation, and on the other hand, asymmetric cell-fates that enable both self-renewal and the generation of new cell types, a single totipotent embryonic cell and its progeny can generate the populations of specified cells that form the diverse tissues and organs of the animal body. In concert with this regulated cellular proliferatiin, gene regulatory networks control the levels and spatiotemporal expression patterns of developmental genes so that proper cell-fates are acquired at the right time and place during development.

*C. elegans* develops through four larval stages (L1-L4). Each larval stage is comprised of an invariant set of cell division and cell fate specification events [1]. The order of cell-fates and the timing of cell-fate transitions within individual cell lineages are regulated by genes in the heterochronic pathway [2,3]. In this pathway, three major temporal regulatory transcription factors control the transitions from earlier to later cell fates. These transcription factors are directly or indirectly regulated by microRNAs and/or RNA binding proteins, thereby facilitating proper cell-fate transitions during larval development.

One of the transcription factors in the heterochronic pathway, Hunchback-like-1 (HBL-1), promotes L2-stage symmetric cell divisions and prevents progression to L3-stage asymmetric cell divisions [4,5]. HBL-1 is expressed at the L1 and L2 stages and is downregulated during the L2-to-L3 transition [4,6]. Proper temporal regulation of HBL-1 activity is characterized by specification of L2 cell-fates at the L2 stage and progression to L3 cell-fates concomitant with the L2-to-L3 stage progression. Mutations that reduce HBL-1 activity during the L1 and L2 stages result in the skipping of L2 cell-fates, whereas mutations that cause ectopic HBL-1 activity at the L3 and L4 stages lead to reiterations of L2 cell-fates at these later larval stages. During the L2-to-L3 transition HBL-1 is regulated by *let-7*-family microRNAs (*mir-48/84/241*) [7] as well as by *lin-28* [7,8], which acts on *hbl-1* indirectly via a protein coding gene *lin-46* [9,10].

LIN-28 is a conserved RNA-binding protein first identified as a heterochronic gene product in *C. elegans* [2]. In *C. elegans* larvae lacking *lin-28* activity, hypodermal stem cells skip L2-stage specific symmetric cell divisions and precociously transition to later stage cell-fates leading to premature terminal differentiation of the hypodermis [2]. LIN-28 inhibits the maturation of the conserved microRNA *let-7* [11], which is required for this hypodermal differentiation at the end of the L4 stage [12]. Curiously, although *let-7* is expressed precociously at the early stages in *lin-28(0)* larvae, and *let-7* function is required for the precocious terminal differentiation of hypodermal cells in *lin-28(0)* animals, *let-7* is not required for the skipping of L2 stage proliferative cell fates observed in *lin-28(0)* [8]. Instead, loss of a protein coding gene, *lin-46*, suppresses both early and late stage *lin-28(0)* phenotypes [9] without repressing precocious *let-7* expression [8]. This suggests that LIN-46 might be mis-regulated in *lin-28(0)* animals and could be responsible for the L2 stage *lin-28(0)* phenotypes irrespective of *let-7*.

Previous studies shown that LIN-28 binds to the *lin-46* mRNA [13] but the consequence of this binding is not known. Additionally, *lin-46* encodes a protein related to a bacterial molybdenum cofactor biosynthesis enzyme and the mammalian protein Gephyrin [14,15], and although the molecular functions of LIN-46 are not clear, our recent findings suggest that LIN-46 affects temporal cell-fates by inhibiting the nuclear accumulation of the transcription factor HBL-1 [10]. Therefore, it is possible that LIN-28 could promote L2 fates by restricting the expression of LIN-46, thereby preserving the nuclear activity of the L2-fate-promoting factor HBL-1.

*C. elegans’* LIN-28 and its homologs in mammals (Lin28) have conserved functions: LIN-28/Lin28 inhibit *let-7* expression [11,16–18], bind to and regulate mRNAs [13,19–21], and promote proliferation and pluripotency [22–24]. Similar to *C. elegans* development, during mammalian embryogenesis LIN-28 is expressed at early/pluripotent stages and is downregulated at later/differentiated stages [22,25,26]. LIN-28 down-regulation in differentiating tissues allows *let-7* microRNA to accumulate resulting in further *let-7*-driven cellular differentiation through the repression of pluripotency and self-renewal promoting genes [27], including *lin-28*. Lastly, LIN-28 expression is associated with many types of cancers and poor prognosis [28,29], and conversely, *let-7* is known to act as a tumor suppressor by repressing oncogenes [30–33].

While certain phenotypes observed in Lin28-deficient mammalian cells can be attributed to increased *let-7* expression and consequent repression of *let-7* targets [18,34], there are *let-7* independent functions of Lin28 [8,35,36], some of which could be explained by mis-regulation of specific mRNA targets of Lin28 [37–40]. Lin28 can regulate the translation of target mRNAs either positively [21] or negatively [19], perhaps in a cell-type specific manner. LIN-28 tends to bind to multiple sites on its mRNA targets [19]. However, the contribution of individual binding sites to mRNA regulation is unclear. Moreover, while the LIN-28-bound mRNA regions are enriched for certain motifs (e.g. GGAG), these sequence motifs are neither required nor sufficient for LIN-28 binding to its targets [19,21]. In brief, the rules and consequences of LIN-28/lin-28 binding to mRNAs are not well understood.

Here we show that the critical target of LIN-28 in *C. elegans*, LIN-46, is expressed only at the L3 and L4 stages in a temporal profile that is the inverse of LIN-28, which is expressed at the L1 and L2 stages. We find that LIN-46 is expressed precociously at the L1 and L2 stages in *lin-28(0)* animals, supporting the idea that LIN-28 represses LIN-46 expression at these early larval stages. We also find that, similar to *lin-28(0)*, mutations in the 5’UTR of *lin-46* result in precocious LIN-46 expression in the hypodermal seam cells, and that this ectopic LIN-46 expression at the L1 and L2 stages is sufficient to result in skipping of L2 stage symmetric seam cell divisions and precocious expression of L3-adult fates. Endogenously tagged LIN-46 is also expressed in the vulval precursor cells (VPCs) and LIN-46 is precociously expressed in the VPCs both in *lin-28(0)* and *lin-46* 5’UTR mutants. Ectopic LIN-46 expression in the VPCs in *lin-46* 5’UTR mutants is sufficient to accelerate cell-fate transitions in these cells, which results in protruding vulva phenotypes similar to *lin-28(0)* animals. Due to the phenotypic similarity between *lin-28(0)* and *lin-46* 5’UTR mutants, we propose that the *lin-46* 5’UTR mediates LIN-28 binding to the *lin-46* mRNA, which results in the repression of LIN-46 expression from the *lin-46* mRNA. Our results demonstrate that precocious LIN-46 expression alone, which is observed in *lin-28(lf)* animals and is sufficient to suppress L2 cell-fates and to induce precocious transition to L3 cell-fates, can account for majority of the *lin-28(lf)* phenotypes.

## RESULTS

### *lin-28* represses LIN-46 expression during early larval stages

The developmental expression patterns of *lin-28* and *lin-46* were previously identified using transcriptional reporters (transgenes expressing a fluorescent protein driven by the promoter of interest) and translational reporters (transgenes expressing the open reading frame of a gene of interest fused with a fluorescent protein driven by the promoter of the gene) [9,22]. The expression of such transgenes does not necessarily represent and accurate readout of expression and spatiotemporal patterns of the endogenous genes. To more accurately determine the expression patterns of LIN-28 and LIN-46, we used CRISPR/Cas9 genome editing to tag *lin-28* and *lin-46* with fluorescent proteins at their endogenous loci (figure 1A). Phenotypic analyses determined that both of the endogenously tagged loci were fully functional. We found that expression pattern of endogenously tagged LIN-28::GFP is comparable to a previous report [22]: LIN-28 is highly expressed in the embryos and at the L1 and L2 stage larvae and is diminished at subsequent stages (figures 1B and 1C). The expression pattern of endogenously tagged LIN-46::mCherry differs from the pattern previously reported for a *lin-46* transcriptional reporter [9]: We observed LIN-46::mCherry predominately in hypodermal seam cells and in the ventral hypodermal vulval precursor cells at the L3 and L4 stages (figure 1C and table S1). Our finding, showing that LIN-46::mCherry expression is restricted to the L3 and L4 stages, combined with a previous report indicating that *lin-46* transcription occurs at all larval stages [9], suggests that *lin-46* is developmentally regulated on the post-transcriptional level.

**Figure 1.**
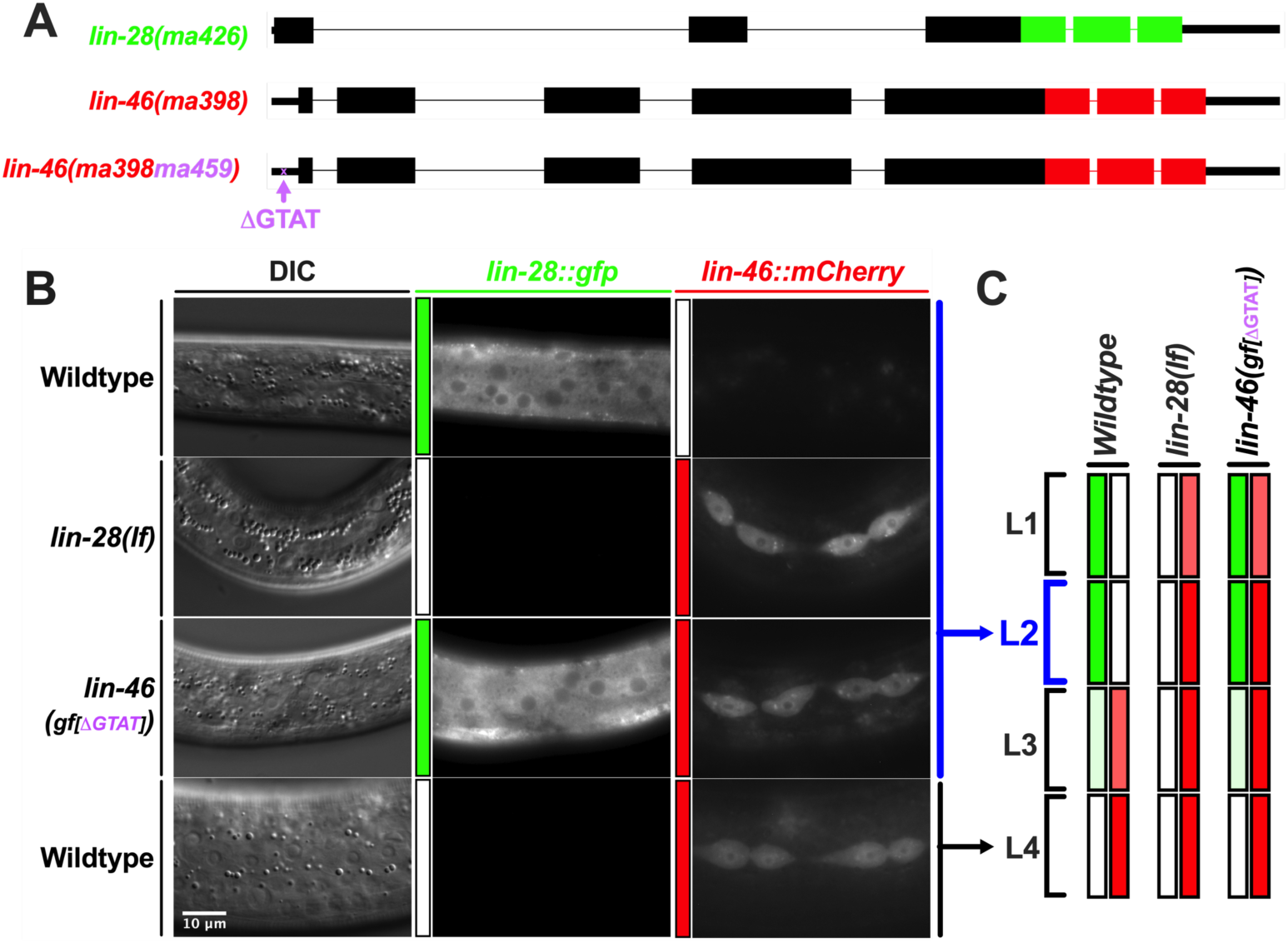
*lin-28* and *lin-46* 5’UTR prevent LIN-46 expression at the L1 and L2 stages. A) Schematic views of the *C. elegans lin-28* and *lin-46* genes and the CRISPR-mediated integration of GFP (green) or mCherry (red) coding sequences at the C-termini. Thick bars represent the exons, thin bars represent the 5’UTRs (left) and 3’UTRs (right). Lines between the exons represent the introns. *Ma426* denotes the endogenously GFP-tagged *lin-28* allele and *ma398* denotes the endogenously mCherry-tagged *lin-46* allele. The *ma398ma459* allele harbors the “GTAT” deletion in the 5’UTR of *lin-46* that is also tagged with mCherry at the C-terminus. B) DIC and fluorescent images showing LIN-28 and LIN-46 expression in wildtype L2 (first row) and L4 (last row) stage larvae, and in *lin-28(lf)* and *lin-46(gf/[*Δ*GTAT])* L2 stage larvae. Fluorescent images of different larvae are taken using the same microscopy setting. All images are then stitched together using the ImageJ software to adjust the brightness and contrast uniformly across the images for enhanced visualization. C) Schematic representation of the LIN-28 and LIN-46 expression observed during L1-L4 larval stages of wildtype, *lin-28(lf)*, and *lin-46(gf/[*Δ*GTAT])* animals. Note that LIN-46 is precociously expressed at the L1&L2 stages in both *lin-28(lf)* and *lin-46(gf[*Δ*GTAT])* mutants.

The expression pattern of endogenously tagged LIN-46 reveals that LIN-28 and LIN-46 are expressed in a temporally mutually exclusive manner: LIN-28 is expressed early (L1 and L2 stages) and LIN-46 is expressed late (L3 and L4 stages). This mutually exclusive expression pattern suggested that LIN-28 may repress LIN-46 expression during the L1 and L2 larval stages. To test this, we examined the effect of loss of *lin-28* on the expression pattern of LIN-46. We found that LIN-46 is expressed precociously at the L1 and L2 stages in *lin-28(0)* animals (figures 1B and 1C), consistent with the conclusion that LIN-28 represses LIN-46 expression at these early larval stages.

### Mutations in the *lin-46* 5’UTR result in *lin-28(0)-*like phenotypes and precocious LIN-46 expression

The finding that LIN-46 is expressed precociously at the L1 and L2 stages in *lin-28(0)* animals suggests two non-mutually exclusive hypotheses: 1) that LIN-28 might directly repress LIN-46 expression, and 2) that precocious LIN-46 expression could contribute to the precocious developmental phenotypes of *lin-28(0)*. The latter hypothesis is supported by previous findings that *lin-46(0)* suppresses the precocious development of *lin-28(0)* [9]. However, while *lin-46* activity was shown to be necessary for the precocious phenotypes of *lin-28(0)*, it is not known if LIN-46 expression is sufficient for precocious development. The possibility that LIN-28 could directly repressed LIN-46 expression by binding to the *lin-46* mRNA in *C. elegans* is supported a previous study showing that LIN-28 could be crosslinked to *lin-46* mRNA in vivo [13].

We sought to test the above hypotheses by mutation of putative LIN-28-interacting sequences in the *lin-46* mRNA sequence and assaying for precocious developmental phenotypes. While the previous study indicated that LIN-28 could be crosslinked to both the 5’ and 3’ UTRs as well as all five exons of the *lin-46* mRNA [13], we noticed that the *lin-46* 5’UTR exhibits unusually high sequence conservation among nematodes (figure S1A) and contains a GGAG motif that is often associated with LIN-28 binding (figures S1A and S1B). Therefore, we targeted the *lin-46* 5’UTR using a CRISPR guide (figures S1A and S1B) and observed frequent *lin-28(0)-*like phenotypes in the F1/F2 progeny of the injected P0 animals (figure S1C). We genotyped several of these F1/F2 progeny, and we found a range of *lin-46* 5’UTR deletions varying in size (2-19 bp) in animals expressing *lin-28(0)-*like phenotypes (Figure S1D).

To determine the effects of the *lin-46* 5’UTR mutations on the expression of LIN-46 we injected the CRISPR mix containing a guide targeting the *lin-46* 5’UTR into animals carrying the *lin-28::gfp [lin-28(ma426)]* and *lin-46::mCherry [lin-46(ma398)]* alleles (figure 1A), and generated *lin-46* 5’UTR deletion mutations, which resulted in precocious LIN-46::mCherry expression at the L1 and L2 stages (figures 1A-1C). This result shows that an intact *lin-46* 5’UTR is required for proper LIN-46 expression. Importantly, our CRISPR mutagenesis of the *lin-46* 5’ UTR did not affect the expression of LIN-28 (figures 1B and 1C). thus LIN-46 is expressed precociously in *lin-46* 5’UTR mutants despite the presence of LIN-28, indicating that the *lin-46* 5’UTR likely mediates LIN-28 binding to, and hence the repression of, the *lin-46* mRNA.

*lin-46* mRNA is expressed at the L1 and L2 stages, fluctuating within a range of approximately two-fold in wild-type animals (figure S2, black line). In *lin-28(0)* and *lin-46* 5’UTR mutants, the *lin-46* mRNA is also expressed at the L1 and L2 stages at similar levels to wild-type. Although at certain time points, *lin-46* mRNA level is slightly but statistically significantly higher or lower in the mutants (figure S2, red and blue lines), it should be noted that these fluctuations in the mRNA levels in wild-type or mutant animals do not correlate with the LIN-46 protein accumulation pattern (figures 1B and 1C); in particular, LIN-46 is never detected at the L1 and L2 stages in wild-type animals, whereas LIN-46 is always present throughout the late L1 and L2 stages in *lin-28(0)* and *lin-46* 5’UTR mutants. Therefore, the precocious LIN-46 expression in *lin-28(0)* and *lin-46* 5’UTR mutants are likely due to an increase in the translatability of the *lin-46* mRNA in these mutants rather than an increase in the *lin-46* mRNA level.

### Precocious LIN-46 expression causes precocious cell-fate transitions in hypodermal seam and vulval cell lineages

To compare in detail the heterochronic phenotypes of three different *lin-46* 5’UTR mutants, *ma461, ma467*, and *ma472*, that are deleted for six, 12, and 19 nucleotides, respectively to those of *lin-28(0)* mutants, we assessed the number of seam cells, the timing of *Pcol-19*::*gfp* expression–a reporter for terminal differentiation of hypodermal cells, the timing of the formation of an adult-specific cuticle structure called alae, and the penetrance of the protruding vulva (Pvl) phenotype (figure 2).

**Figure 2.**
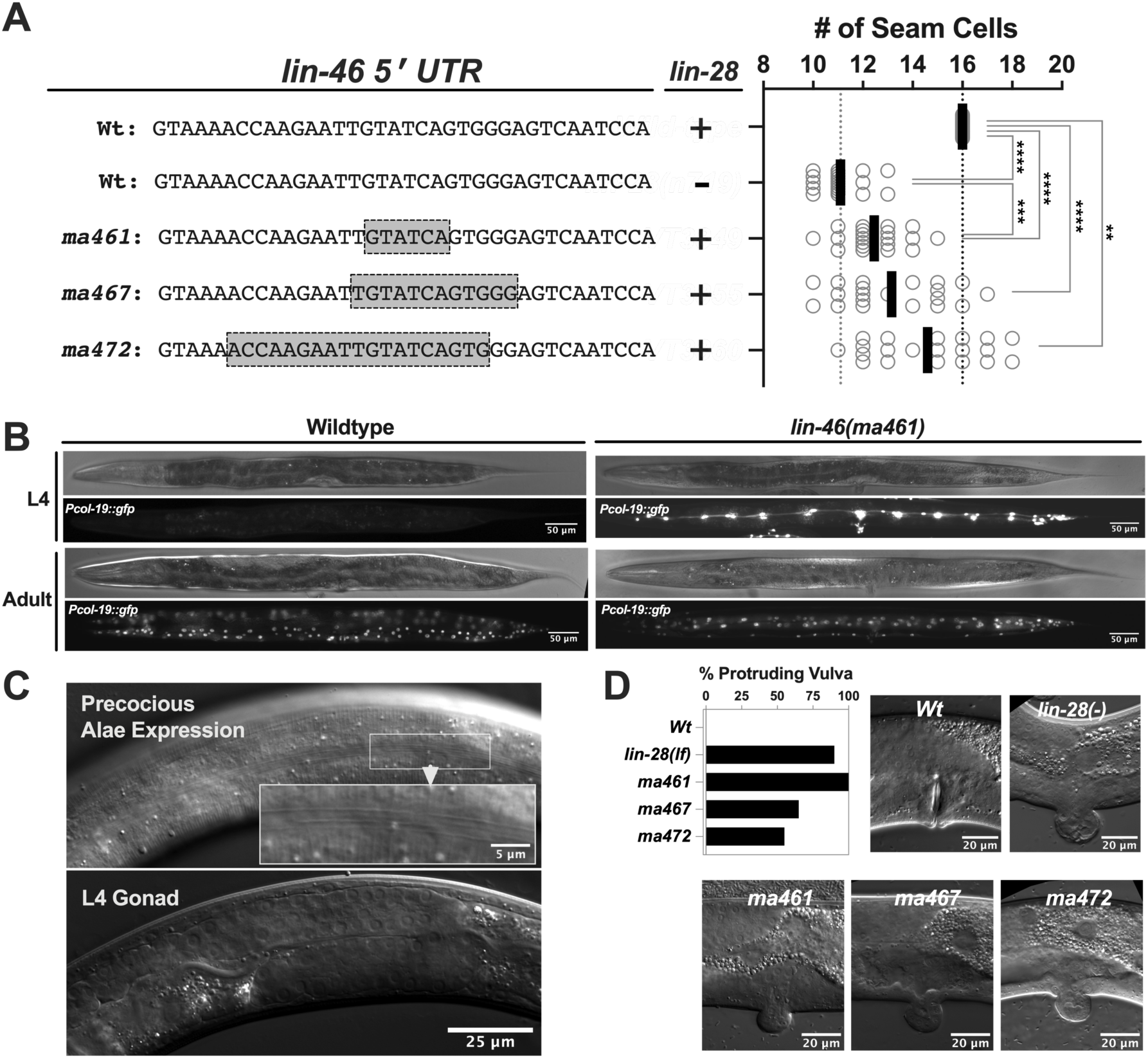
Precocious LIN-46 expression causes precocious cell-fate transitions in hypodermal seam and vulval cell lineages. A) Number of seam cells observed in young adults of wildtype, *lin-28(lf)*, and three *lin-*46 5’UTR mutants (*ma461, ma467, and ma472*). Each dot in the plot on the right represents the number of seam cells observed on one lateral side of a single worm. Vertical black bars indicate the average number of seam cells observed for each genotype. B) DIC and fluorescent images of L4 and adult stage wildtype and *lin-46(ma461*) animals. The adult specific *Pcol-19::gfp (maIs105)* is normally expressed in the hypodermal seam and hyp7 cells at the adult stage of both wildtype and *lin-46(ma461)* animals. However, unlike in wildtype larvae, *Pcol-19::gfp* is also precociously expressed in the seam cells of L4 stage *lin-46(ma461)* larvae. C) DIC images showing the (precocious) adult-specific cuticle structure called alae (upper panel) on the cuticle of a larva at the L4 stage indicted by the developmental stage of the gonad (lower panel). D) Percent protruding vulva (Pvl) phenotype observed in wildtype, *lin-28(lf)*, and three *lin-46* 5’UTR mutants and DIC images showing normal vulva or Pvl morphology observed in each genotype.

First, all three *lin-46* 5’UTR mutants had fewer seam cells than wildtype (figure 2A), which indicates that *lin-46* 5’UTR mutations, presumably as a consequence of consequent precocious LIN-46 expression, skip the L2-stage symmetric seam cell divisions. The severity of this seam cell phenotype varies for each mutant and is weaker than *lin-28(0)* (figure 2A). The variability in the number of seam cells is due to variation in cell-fate decisions across the seam cells of each larva; namely, in *lin-28(0)* animals almost all seam cells skip L2 cell-fates whereas in the 5’UTR mutants of *lin-46* not all but only some (and a varying number of) seam cells skip L2 stage cell-fates. It also worth noting that while the majority of the *ma472* animals had fewer than sixteen seam cells, some *ma472* animals had more than sixteen seam cells (figure 2A) indicating that at least certain seam cells reiterated L2 cell-fates at later stages, which might be due to reduced *lin-46* expression in those seam cells. This variation in the number of seam cells that skip or reiterate L2 fates in *lin-46* 5’UTR mutants could reflect a variability in the timing or level of LIN-46 expression across seam cells due to the misregulation of *lin-46* translation.

Second, similar to what is observed in *lin-28(0)* animals, seam cells in *lin-46* 5’UTR mutants precociously express adult fates during larval stages as demonstrated by the precocious expression of *Pcol-19::gfp*, (figure 2B) and by the expression of an adult alae in L4 stage larvae (figure 2C).

Lastly, we quantified the percent animals that display protruding vulva (Pvl) phenotypes in young adults (a characteristic of precocious vulval development in *lin-28(0)* animals): all three *lin-46* 5’UTR mutants displayed Pvl phenotypes similar to *lin-28(0)* animals (figure 2D).

### LIN-46 is expressed in the vulval precursor cells (VPCs) and precocious LIN-46 expression leads to precocious onset of vulva development

During *C. elegans* larval development, stem cells of the ventral hypodermal lineages P3-P8 divide during the L1 stage and give rise to the six P3.p-P8.p VPCs [1]. After their birth in the L1 stage, the VPCs temporarily arrest in the G1 phase of the cell cycle until the L3 stage when they undergo a single round of cell division (figure 3A) [41]. Concomitant with this cell division, three of the six VPCs (P5.p, P6.p, and P7.p) undergo additional rounds of cell divisions, giving rise to 22 cells that progressively differentiate and form the adult vulva (figure 3A). The timing of the first VPC divisions in the mid-L3 stage is controlled by genes in the heterochronic pathway, including *lin-28* [2,41]. In *lin-28(0)* mutants, the first VPC divisions precociously take place in the L2 stage; the VPC progeny subsequently continue to precociously divide and differentiate, resulting in precocious vulva development, evidenced by an abnormally formed, protruding vulva. Loss of *lin-46* suppresses the Pvl phenotype caused by *lin-28(0)*. However, because LIN-46 expression in the VPCs had not been previously detected using transgenic reporters [9], it was not clear how loss of *lin-46* could affect the Pvl phenotype of *lin-28(0)* animals.

**Figure 3.**
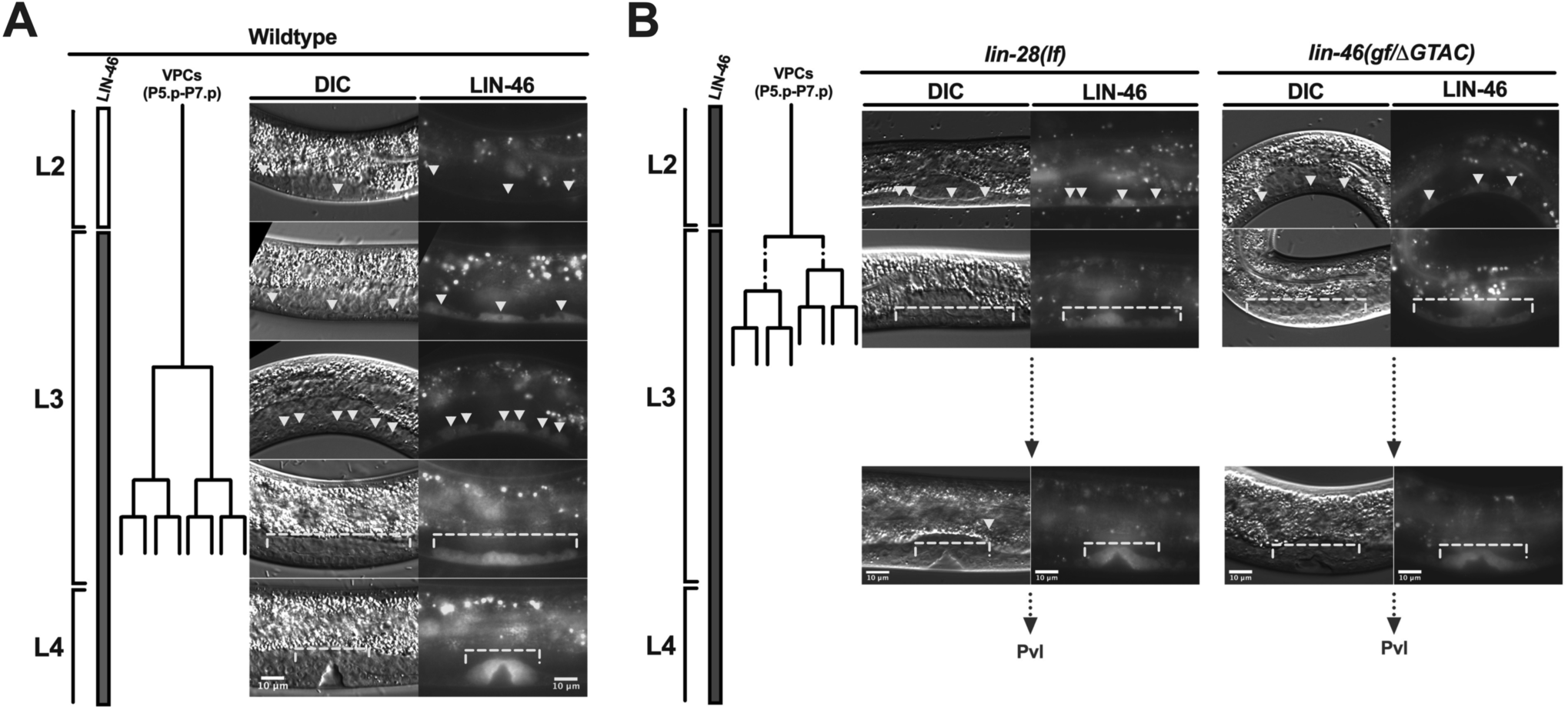
LIN-46 is expressed in the vulval precursor cells (VPCs) and precocious LIN-46 expression leads to precocious onset of vulva development. A) Wildtype larval stages and the cell lineage diagram of the vulval precursor cells (VPC), and DIC and fluorescent showing LIN-46 expression in the VCPs (indicated by arrowheads or brackets). B) Larval stages (according to gonad morphology), precocious LIN-46 expression and VPC development in *lin-28(lf)* and *lin-46(gf[*Δ*GTAT])* animals are shown. Importantly, the synchrony among P5.p, P6.p and P7.p and among their progeny are lost in *lin-28(lf)* and *lin-46(gf[*Δ*GTAT])*. Accordingly, due to the variability (indicated by dotted lines in the lineage) in the timing of the VPC divisions, the distinct temporal patterns of vulva development observed in the wildtype animals from early to late L3 stage (as in Panel A) is also lost.

We found that endogenously tagged LIN-46 is expressed in the VPCs at the L3 and L4 stages (figure 3A), which coincides with the period when the VPCs develop into adult vulva. In *lin-28(0)* mutants, LIN-46 is precociously expressed in the L2-stage VPCs, which coincides with its precocious VPC development (figure 3B, left). Similarily, in *lin-46* 5’UTR mutants, LIN-46 is precociously expressed in the VPCs at the L2 stage coinciding with their VPCs precocious development in these mutants (figure 3B, right). This indicates that precocious LIN-46 expression is sufficient to alter the timing of vulva development inferring that precocious LIN-46 expression is likely responsible for precocious vulva development in *lin-28(0)* animals. Altogether, our results show that the LIN-28 target LIN-46 suppresses L2 cell-fates to promote the transition to L3 cell-fates in both seam cells and vulval precursor cells, and that precocious LIN-46 expression causes precocious cell-fate transitions, which is likely responsible for the majority of the phenotypes observed in *lin-28(0)* animals.

### The architecture and conservation of the *lin-46* 5’UTR

Most *C. elegans* transcripts are trans-spliced resulting in the fusion of a 22-nt splice-leader (SL) RNA to the 5’ end of the transcripts [42]. The presence of an upstream splice acceptor “TTTCAG” (figures 4 and S1B) and expressed sequence tag (EST) clones that contain the *SL1*-*lin-46*-5’UTR fusion sequence (e. g. GenBank: FN875625.1) indicate that *lin-46* mRNA is trans-spliced. We used the RNAfold Webserver [43] to predict the structure of the *SL1*-*lin-46*-5’UTR fusion sequence (figure 4A and figure S3A). The predicted structure of the *SL1*-*lin-46*-5’UTR chimeric RNA shows base-pairing between *SL1* and the first eight nucleotides at the 5’ end of the *lin-46* 5’UTR (figure 4A, “*SL1*-complementary”). These eight nucleotides are highly conserved across *Caenorhabditis* species (figure 4B); and a nucleotide variation at position four that is found in five different species preserve the predicted base-pairing (A-U to G-U) with the *SL1* sequence, which supports the biological relevance of the predicted structure (figure 4). Perturbing this *SL1*-complementary region in addition to mutating more downstream sequences in the 5’UTR resulted in a weaker phenotype (*ma461* versus *ma472* in figure 2A, D, S3B), which suggests that these first eight nucleotides or their base-pairing with the *SL1* sequences has a positive impact on LIN-46 expression.

**Figure 4.**
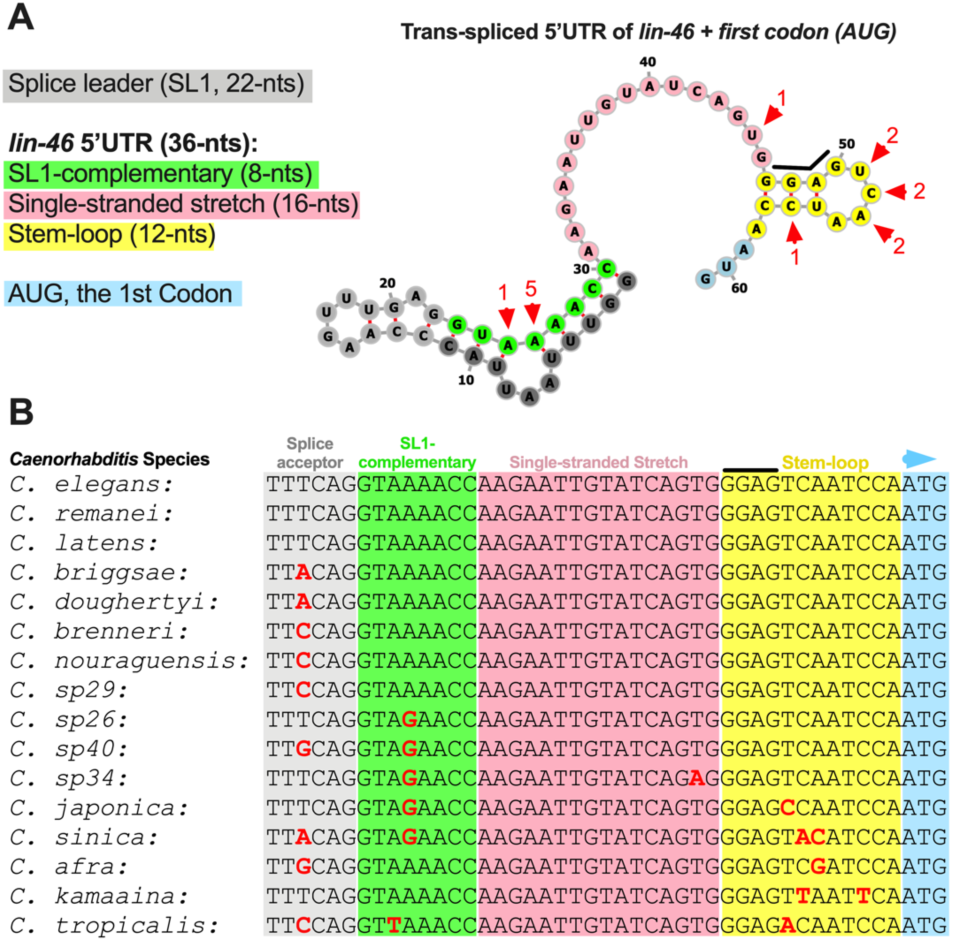
The architecture and conservation of the *lin-46* 5’UTR. A) The predicted structure of the trans-spliced *lin-46* 5’UTR and the annotation of distinct structural regions. Red arrowheads indicate nucleotides that are mutated in nematode species (as listed in panel B); numbers denote the number of times a mutation is found at the position indicated. B) An alignment of the genomic sequences encoding the *lin-46* 5’UTR in various *Caenorhabditis* species. The *lin-46* 5’UTR is highly conserved and the conservation pattern is consistent with the conservation of the base-pairing interactions in the predicted structure: base-pairing nucleotides are more conserved or nucleotide changes preserve base-pairing (e.g. A:U to G:U).

Sixteen nucleotides that follow the *SL1*-complementary region are conserved among all *Caenorhabditis* species analyzed here and constitute a “single-stranded stretch” region (figure 4A) with the exception of *C. sp34* that has a single nucleotide change in this region (figure 4B). This single-stranded stretch was the region primarily targeted by our CRISPR guide used to generate the 5’UTR mutants (figure S1B). Mutations of various sizes (figure S1D) in this region alone displayed precocious LIN-46 expression (figure 1A-C, *ma459*) and strong *lin-28(0)*-like phenotypes (figure 2A, *ma461*). Interestingly, in certain *lin-46* 5’UTR mutants, such as the *ma459*, that result in strong precocious phenotypes, the predicted RNA structure is entirely altered (figure S2B), which may indicate a causative relationship between loss of all structural elements in the *lin-46* 5’UTR and a strong loss of LIN-46 repression.

The last 12 nucleotides in the *lin-46* 5’UTR contains a GGAG sequence that is located in the stem of a predicted stem-loop structure (figure 4A, stem-loop). The sequence conservation pattern in this region supports the biological relevance and significance of the predicted structure: 1) a C to T nucleotide change in the stem preserves base-pairing (G-C to G-U), and 2) mutations in the nucleotides in the loop region, which are not contributing to the hairpin stability, appear to be more tolerated (figure 4). The GGAG motif is found to be enriched in LIN-28 bound RNA regions [13], however, here the GGAG sequence or the stem-loop in the *lin-46* 5’UTR alone is not sufficient to confer repression of LIN-46 expression (see *ma472* in figure 2A and figure S2). Moreover, perturbing the GGAG sequence in addition to the single-stranded stretch sequences did not enhance but rather weakened the precocious phenotypes (*ma467* vs *ma461* in figures 2A, D and S3), which suggests that this GGAG-containing loop, rather than an having an inhibitory role, can positively affect LIN-46 expression.

## DISCUSSION

Our results provide insights into how the conserved RNA-binding protein LIN-28 regulates its critical mRNA target, *lin-46*, in *C. elegans*, and demonstrate that *lin-46* mis-regulation is likely responsible for the phenotypes observed in *lin-28(0)* animals. Our results suggest that LIN-28 controls temporal cell-fate progression by regulating LIN-46 expression via the 5’UTR of *lin-46* mRNA.

The temporally mutually exclusive expression pattern between LIN-28 and LIN-46 (revealed by the endogenously tagged alleles of *lin-28* and *lin-46*) and the effect of loss-of-function of *lin-28* on the LIN-46 expression led us to conclude that *lin-28* represses LIN-46 expression at early stages (figure 5). Our measurements of *lin-46* mRNA levels in lin-28 mutants indicated that LIN-28 likely regulates LIN-46 production at the level of translation. Our results also suggest that the 5’UTR of *lin-46* prevents LIN-46 expression, which is likely via mediating LIN-28 binding to and repression of the *lin-46* mRNA. We showed that LIN-46 is precociously expressed in *lin-28(0)* animals; with the help of the *lin-46* 5’UTR mutants, which uncouple precocious LIN-46 expression from the loss-of-function of *lin-28*, we showed that precocious LIN-46 expression alone is sufficient to suppress L2 cell-fates and to promote precocious transitions to L3 cell-fates (figure 2). Lastly, endogenously tagged LIN-46 is expressed in the VPCs, which had not been reported before, and *lin-28* and portions of the *lin-46* 5’UTR are required to repress LIN-46 expression in the VPCs. Precocious onset of LIN-46 expression in the VPCs is sufficient to stimulate the precocious onset of vulva development (figure 3). These results demonstrate that precocious LIN-46 expression in seam cells and VPCs in *lin-28(0)* mutants is responsible for the two major heterochronic phenotypes observed in the *lin-28(0)* animals; skipping of L2 stage seam cell proliferation and precocious onset of vulva development.

**Figure 5.**
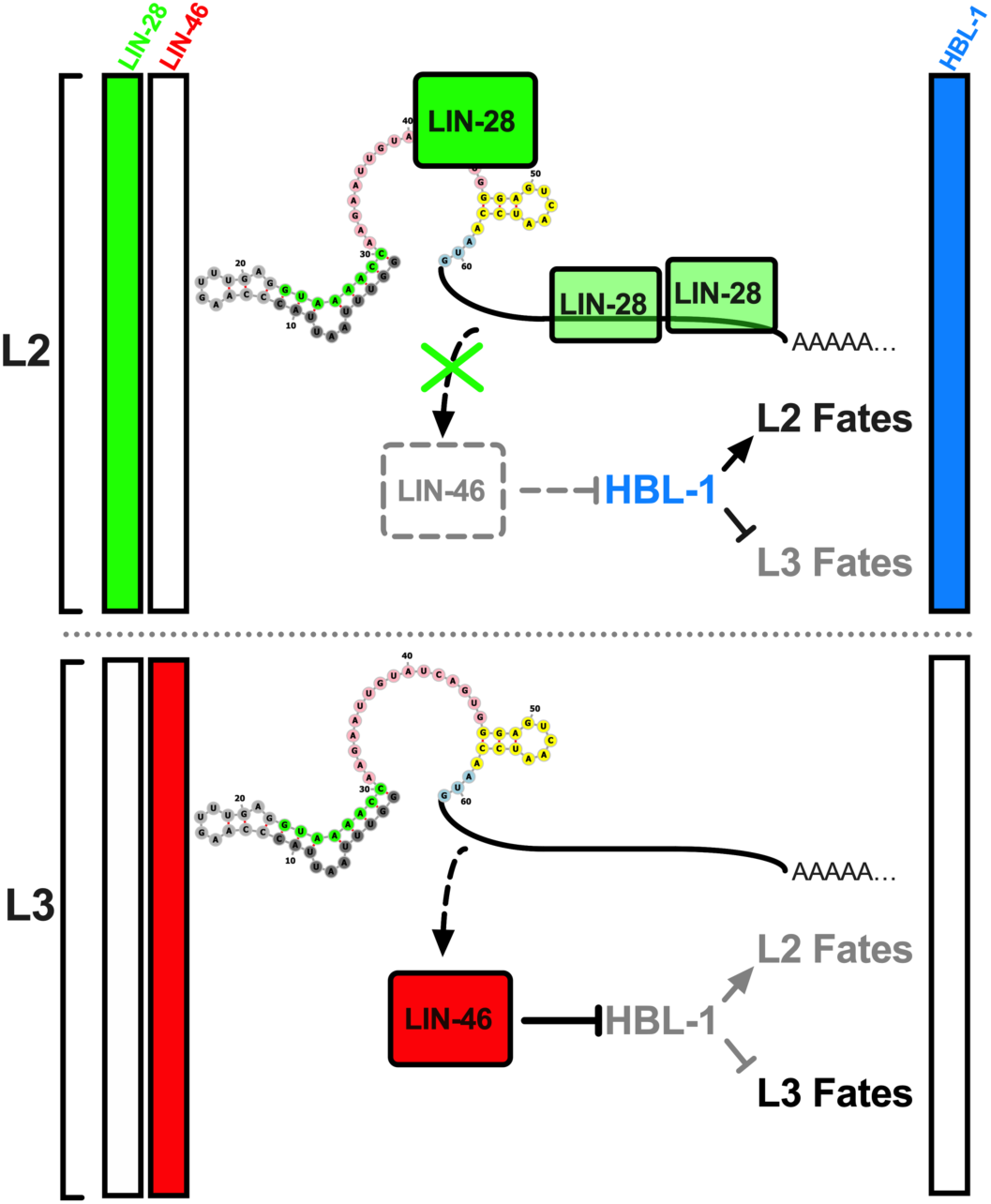
Model: LIN-28 controls temporal cell-fate progression by regulating LIN-46 expression via the 5’UTR of *lin-46* mRNA. At the L2 and L3 stages LIN-28 is highly expressed and although the *lin-46* mRNA is transcribed it cannot be translated due to LIN-28-mediated inhibition of its translation. Inhibition of LIN-46 expression at the L2 stage permits HBL-1 to function: HBL-1 promotes L2 cell-fates and prevents L3 cell-fates. At the L2 to L3 molt, LIN-28 expression is diminished, and this allows LIN-46 expression. LIN-46 opposes HBL-1 activity, thereby preventing expression of L2 cell-fates at the L3 stage.

We hypothesize that the *lin-46* 5’UTR contains a LIN-28-binding element that is required for LIN-28-mediated repression of LIN-46 expression from the *lin-46* mRNA. The evidence that supports this hypothesis include: 1) The phenotypic similarities between *lin-28(0)* and the 5’UTR mutants of *lin-46* reported here; 2) LIN-28 binding to the *lin-46* mRNA (including the *lin-46* 5’UTR) reported previously [13], 3) The existence of a putative LIN-28 interacting sequence, the “GGAG”, in the 5’UTR of *lin-46* (figure 4). However, because previously LIN-28 was shown to interact with the *lin-46* mRNA at multiple sites across the entire *lin-46* mRNA in addition to the 5’UTR [13], it is surprising that mutations of the *lin-46* 5’UTR are sufficient to cause a phenotype that is consistent with an almost total loss of LIN-28-mediated regulation of the *lin-46* mRNA. Nonetheless, at least two models could reconcile a potential total loss of LIN-28-mediated repression of LIN-46 expression by mutating the 5’UTR of *lin-46* alone while leaving other LIN-28 binding sites on the *lin-46* mRNA intact. The first model is that the binding of LIN-28 to the *lin-46* 5’UTR would be required for inhibiting LIN-46 expression whereas LIN-28 binding to other sites on the *lin-46* mRNA would not inhibit LIN-46 expression. It is known that translation initiation is highly regulated [44]; and the 5’UTRs harbor sequence elements, such as upstream open reading frames (uORFs), or structural elements, such as highly structured RNA (including G-quadruplexes and pseudoknots) or specific RNA structures that serve as binding sites for RNA-binding proteins [45], which can interfere with or inhibit the translation initiation [46,47]. In the second model, among all the LIN-28 binding sites on the *lin-46* mRNA, the *lin-46* 5’UTR (and particularly the single-stranded stretch region) might have the highest affinity for LIN-28 and might be required to initiate a sequential binding of multiple LIN-28 proteins to the *lin-46* mRNA, leading to the formation of a repressive LIN-28-*lin-46*-mRNA mRNP (messenger ribonucleoprotein) complex. In support of this model, in *in vitro* assays, LIN28 is shown to preferentially bind to single stranded RNA and more than one LIN28 proteins are shown to bind to long (longer than 30-nts) RNA in a sequential manner after the first LIN28 binds to a predicted single-stranded loop [21].

Precocious cell-fate transition phenotypes observed in *lin-46* 5’UTR mutants are not as strong as the phenotypes observed in *lin-28(0)* mutants. Moreover, the severity of the *lin-28(0)*-like phenotypes vary among different *lin-46* 5’UTR alleles. The severity of these phenotypes does not correlate with the size of the *lin-46* 5’UTR deletions; and in some cases, larger deletions result in not stronger but more moderate phenotypes (figure 2). These findings are consistent with a model where the *lin-46* 5’UTR harbors multiple cis-regulatory elements that can either positively or negatively affect LIN-46 expression. Accordingly, mutants that inactivate a negative regulatory element (the presumed LIN-28 binding site) without disturbing a positive regulatory element result in higher LIN-46 expression and hence stronger precocious phenotypes.

A putative positive regulatory element in *lin-46* 5’UTR could be the first eight nucleotides of the 5’UTR that are predicted to base-pair with the *SL1* sequence (*SL1*-complementary, figure 4). The *lin-46* mRNA is trans-spliced, which results in the fusion of the *SL1* RNA to the 5’ end of the transcript (Figure 4 and S3A). A small stem-loop structure in the *SL1* sequence (Figure S3A) has been shown to enhance translation in nematodes [48]. In the predicted folding of the *SL1*-*lin-46*-5’UTR chimeric RNA (Figure S3A), the nucleotides that form the stem-loop in the *SL1* alone base-pair with the first eight nucleotides of the *lin-46* 5’UTR (Figure S3A). This *SL1*-5’UTR base-pairing is lost in the *lin-46(ma472)* (Figure S3B) and the phenotype of this larger deletion is weaker than the two other, smaller deletions (Figure 2A and S3B), which is consistent with the idea that base-pairing of *SL1* to the *lin-46* 5’UTR has a positive impact on the translatability of the *lin-46* mRNA.

The *C. elegans lin-28-lin-46* pathway acts in parallel to *let-7*-family microRNAs [7,8] and regulates the nuclear localization, hence the activity, of a critical *let-7*-family target and a transcription factor, HBL-1 [10]. *lin-46* activity becomes more important in preventing L3/L4-stage HBL-1 activity at low temperatures [9], or when animals develop through a temporary diapause [49], or merely when animals experience an extended L2 (called L2d) in the presence of diapause-inducing pheromones or starvation stress [6]. L2d/Dauer-inducing conditions also result in the repression of *let-7*-family microRNAs [50,51]. Therefore, LIN-28-mediated regulation of *lin-46* mRNA translation might play a role in controlling a compensation mechanism against environmentally-induced reduction in *let-7*-family levels, perhaps by regulating the level of LIN-46 accumulation at the early L3 stage.

Remarkably, precocious LIN-46 expression conferred by the *lin-46* 5’UTR mutants can fully compensate for the loss of all ten *let-7-*complementary sites in the *hbl-1* 3’UTR [10], which otherwise causes a severe extra seam cell phenotype due to ectopic HBL-1 activity at L3/L4 stages [10]. Therefore, one way to compensate for reduced *let-7*-family levels by LIN-46 activity would simply be the induction of LIN-46 expression in response to environmental conditions that repress *let-7*-family microRNAs. Such an induction mechanism that regulates LIN-46 levels at the L3 stage to match the level of *let-7-*family repression at the L2 stage may utilize LIN-28/5’UTR-mediated regulation of LIN-46 expression as a gate to uncouple *lin-46* mRNA accumulation from LIN-46 protein accumulation, which can provide a control over the rate of LIN-46 accumulation at the early L3 stage. In this hypothetical model, during a lengthened L2 (L2d), *lin-46* mRNA may accumulate in proportion to the length of the L2d stage (which is thought to correlate with the severity of the environmental conditions as well as the degree of *let-7*-family repression). At the L2/L2d stage *lin-46* mRNA cannot be translated due to LIN-28-mediated inhibition; however, at the L3 stage, when LIN-28 expression is diminished, the L2/L2d-accumulated pool of *lin-46* mRNA would be translated. Thus, LIN-46, expressed from a *lin-46* mRNA pool whose size negatively correlates with the reduction in *let-7*-family levels, can accumulate fast enough to sufficiently inhibit the residual or ectopic HBL-1 at the post-L2d L3 stage, compensating for the reduced *let-7*-family microRNA levels by an appropriately matching level of LIN-46.

3’UTR- and microRNA-mediated mechanisms and their roles in controlling temporal dynamics of gene expression have extensively been studied in the context of the *C. elegans* heterochronic pathway. However, the involvement of 5’UTRs in the heterochronic pathway was not known and the identities and roles of cis-regulatory elements in the *C. elegans* 5’UTRs are largely unknown. Conservation in the 5’UTRs is not widespread, but interestingly, the 5’UTRs of many heterochronic and developmental genes in *C. elegans* appear to be evolutionary conserved, which may provide a platform to further explore the functions of mRNA cis-regulatory elements and the roles of trans-acting RNA binding proteins in regulating stage specific gene expression, and developmental progression as well as other important developmental processes.

In summary, we provide evidence indicating that LIN-28 represses the expression of its critical mRNA target in *C. elegans*, an intact *lin-46* 5’UTR is required LIN-28-mediated repression of *lin-46* expression, and precocious LIN-46 expression alone is likely responsible for the majority of *lin-28(lf)* phenotypes. Our findings highlight the biological importance of the mRNA targets of LIN-28 (*C. elegans* LIN-28 and its orthologs), which may have important functions in regulating pluripotency, reprogramming, or oncogenesis in humans and various other organisms.

## MATERIALS AND METHODS

### *C. elegans* culture conditions

*C. elegans* strains used in this study and corresponding figures in the paper are listed in Table S2. *C. elegans* strains were maintained at 20°C on nematode growth media (NGM) and fed with the *E. coli* HB101 strain.

### Assaying extra seam cell and Pvl phenotypes

The worms were scored at the young adult stage (determined by the gonad development) for the number of seam cells using fluorescence microscopy with the help of the *maIs105* [*pCol-19::gfp*] transgene that marks the lateral hypodermal cell nuclei and/or for protruding vulva phenotype (Pvl) by examining the vulva morphology (as given in Figure 2D).

Each circle on the genotype versus number of seam cells plots shows the observed number of seam cells on one side of a single young adult worm. A minimum of 20 worms for each genotype are analyzed and the average number of seam cells (denoted by lateral bars in the genotype versus number of seam cell plots); percent Pvl values are calculated and represented using a bar graph. The Student’s t test is used to calculate statistical significance when comparing different genotypes. The GraphPad Prism 8 software is used to plot the graphs and for statistical analysis.

### Microscopy

All DIC and fluorescent images are obtained using a ZEISS Imager Z1 equipped with ZEISS Axiocam 503 mono camera, and the ZEN Blue software. Prior to imaging, worms were anesthetized with 0.2 mM levamisole in M9 buffer and mounted on 2% agarose pads. The ImageJ Fiji software is used to adjust the brightness and contrast of the images to enhance the visualization of the fluorescent signal. All images are taken using the same microscopy settings and a standard exposure time for all larval stages and genetic background for each reporter (*lin-28::gfp* and *lin-46::mCherry*). To enhance the visualization of the fluorescent signals in the figures and to allow comparison of signal intensities in larvae of different genetic backgrounds, fluorescent images of larvae from different backgrounds are stitched together using the ImageJ software and the brightness and contrast of these montaged images were adjusted (in Figure 1B and 3).

### Tagging of lin-28 and *lin-46* using CRISPR/Cas9

A mixture of plasmids encoding SpCas9 (pOI90, 70 ng/μL), and single guide RNAs (sgRNAs) targeting the site of interest (60 ng/μL of pSW65 for *lin-28* or pOI113 for *lin-46*) and the *unc-22* gene (pOI91, 30 ng/μL) as co-CRISPR marker, a donor plasmid (20 ng/μL of pOI173 for lin-28 or pOI167 for lin-46) containing the gfp or mCherry sequence flanked by gene-specific homology arms, and a *rol-6(su1006)* containing plasmid (pOI124, 30 ng/μL) as co-injection marker was injected into the germlines of ten young adult worms. F1 roller and/or twitcher animals (100-200 worms) were cloned and screened by PCR amplification (Table S3) for the presence of the expected homologous recombination (HR) product. F2 progeny of F1 clones positive for the HR-specific PCR amplification product were screened for homozygous HR edits by PCR amplification of the locus using primers that flanked the HR arms used in the donor plasmid (Table S3). Finally, the genomic locus spanning the HR arms and *gfp* or *mCherry* DNA was sequenced using Sanger sequencing. A single worm with a precise HR edited *lin-28* or *lin-46* locus was cloned and backcrossed twice before used in the experiments.

### CRISPR/Cas9-mutagenesis of the *lin-46* 5’UTR

A mixture of plasmids encoding SpCas9 (pOI90, 70 ng/μL), and gR_5U single guide RNA (sgRNAs) targeting the *lin-46* 5’UTR (Figure S1; pOI193 60 ng/μL) was injected into the germlines of young adult worms expressing the adult onset *gfp* transgene (Table S2; VT1357). F1 or F2 animals displaying precocious cell-fate phenotypes, which were consisted of precocious *Pcol-19::gfp* expression in the seam cells (Figure S2C) and protruding vulva morphology (Figure 2D), were cloned and genotyped for in-del events at the gR_5U targeting site (Figure S2D).

### Quantitative PCR

Samples of total RNA were pre-treated with turbo DNase (Invitrogen). cDNA was synthesized using SuperScript IV (Invitrogen) following the manufacturer’s instructions, using the RT oligonucleotide “oligo (dT)”. qPCR reactions were performed using CoWin Biosciences FastSYBR Mixture (Low Rox) following the manufacturer’s instructions, using a Viia 7 Real Time PCR System (Applied Biosystems). ΔCTs were calculated by normalizing samples to *gpd-1* (*GAPDH*). ΔCTs were then inverted so that greater values reflect greater RNA levels, and were normalized to set the value of the least abundant sample to one. For each biological replicate, the average of three technical replicates was used.

## Supplemental information

**Table S1.**
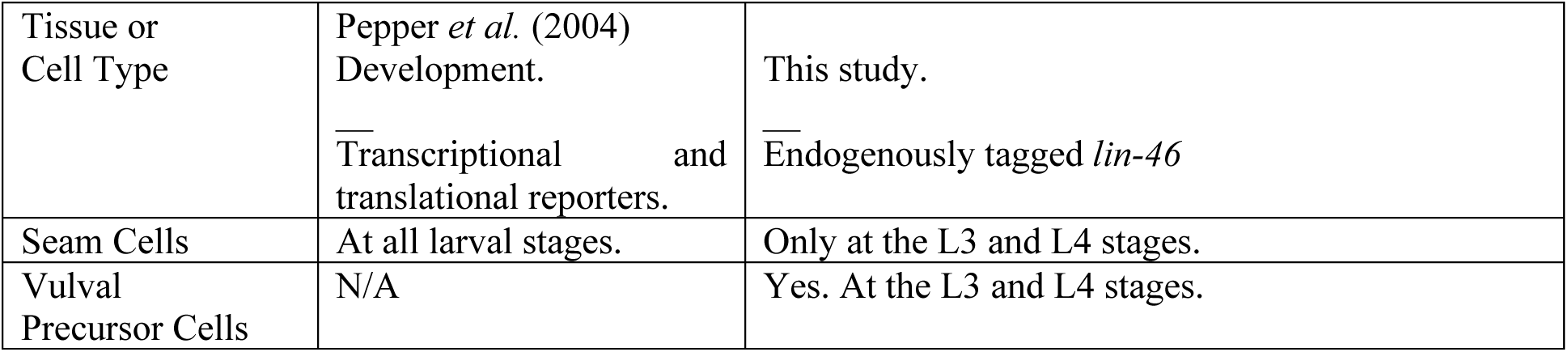
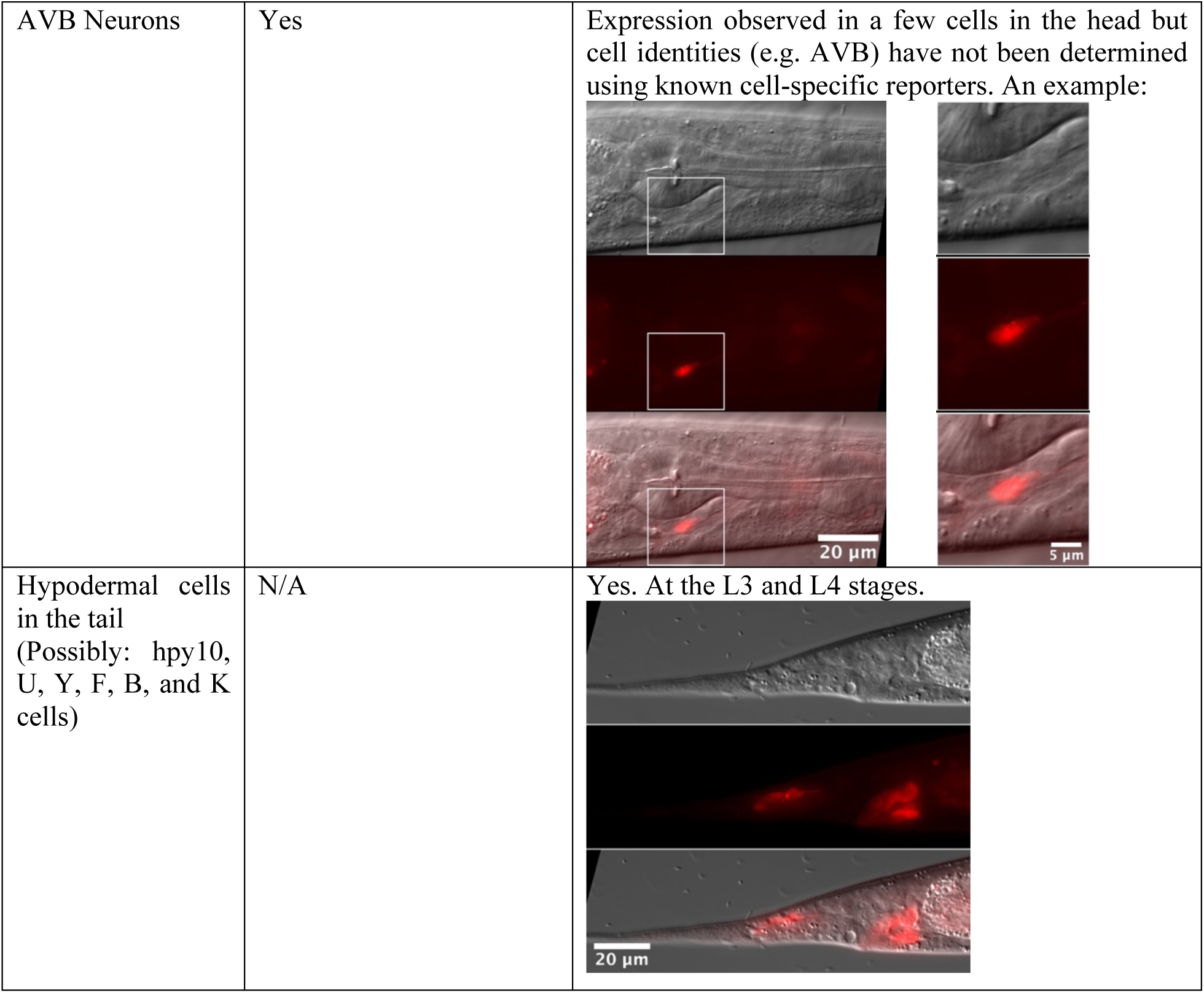
Comparison of LIN-46 expression observed in transgene reporters versus endogenously tagged locus.

| Tissue or Cell Type | Pepper *et al.* (2004)<br>Development.<br>Transcriptional and translational reporters. | This study.<br>Endogenously tagged *lin-46* |
| --- | --- | --- |
| Seam Cells | At all larval stages. | Only at the L3 and L4 stages. |
| Vulval Precursor Cells | N/A | Yes. At the L3 and L4 stages. |
| AVB Neurons | Yes | Expression observed in a few cells in the head but cell identities (e.g. AVB) have not been determined using known cell-specific reporters. An example: |
| Hypodermal cells in the tail<br>(Possibly: hpy10, U, Y, F, B, and K cells) | N/A | Yes. At the L3 and L4 stages. |

**Table S2.** *C. elegans* strains used in this study.

| Strain name | Genotype | Related figures |
| --- | --- | --- |
| VT3737 | *lin-28(ma426[lin-28::gfp] I; lin-46(ma398[lin-46::mCherry] V* | Figure 1A-C, 3A |
| VT3652 | *lin-28(n719) I; lin-46(ma398[lin-46::mCherry] V* | Figure 1A-C, 3B |
| VT3847 | *lin-28(ma426[lin-28::gfp] I; lin-46(ma398ma459[ΔGTAT::lin-46::mCherry] V* | Figure 1A-C, 3B |
| VT1367 | *maIs105 (Pcol-19::gfp) V* | Figure 2A, B, D, S1C |
| VT790 | *lin-28(n719) I; maIs105 V* | Figure 2A, D |
| VT3849 | *lin-46(ma461) maIs105 V* | Figure 2A-D |
| VT3855 | *lin-46(ma467) maIs105 V* | Figure 2A, D |
| VT3860 | *lin-46(ma472) maIs105 V* | Figure 2A, D |

**Table S3.**
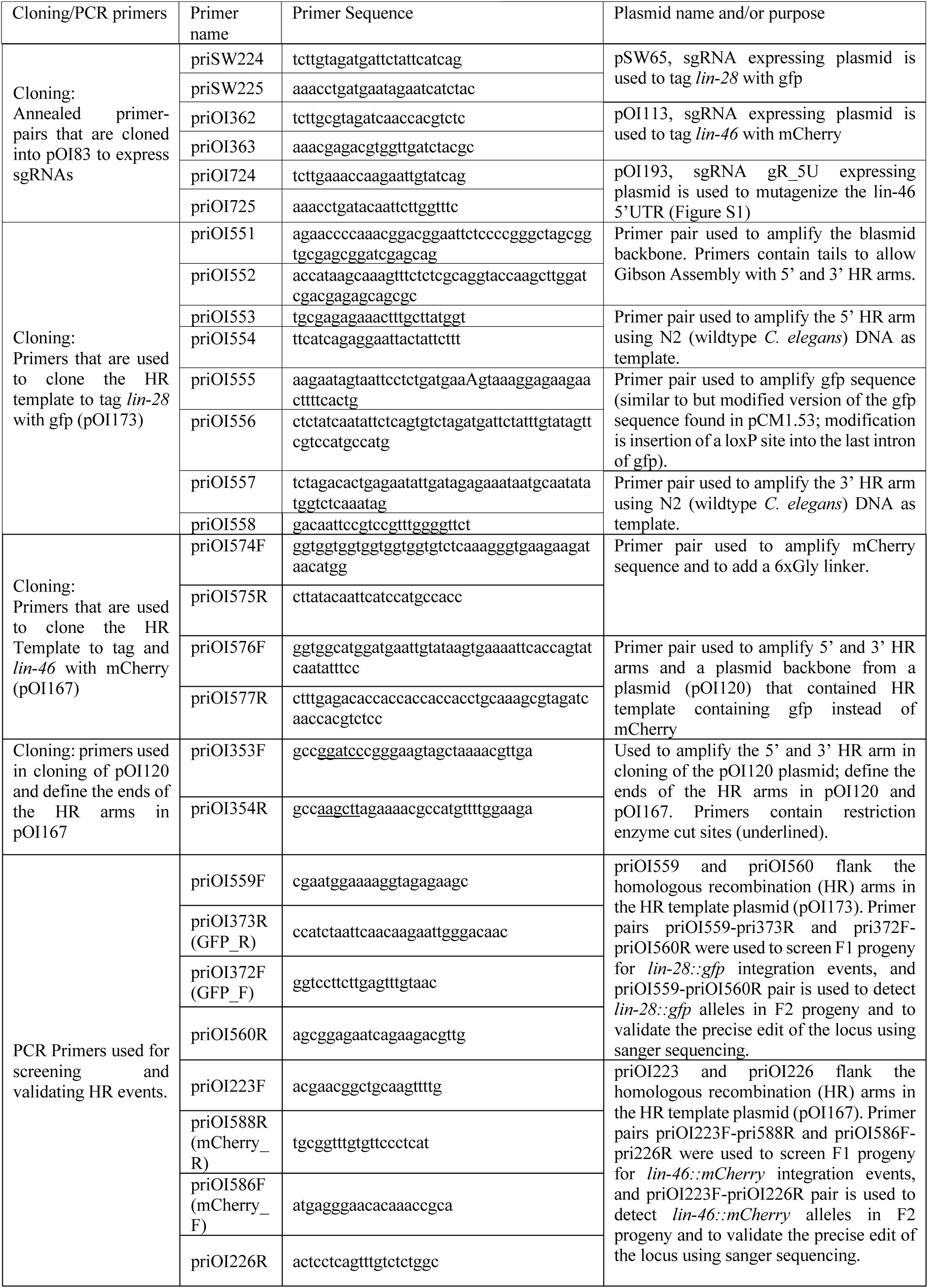
PCR Primers used in this study.

**Figure S1.**
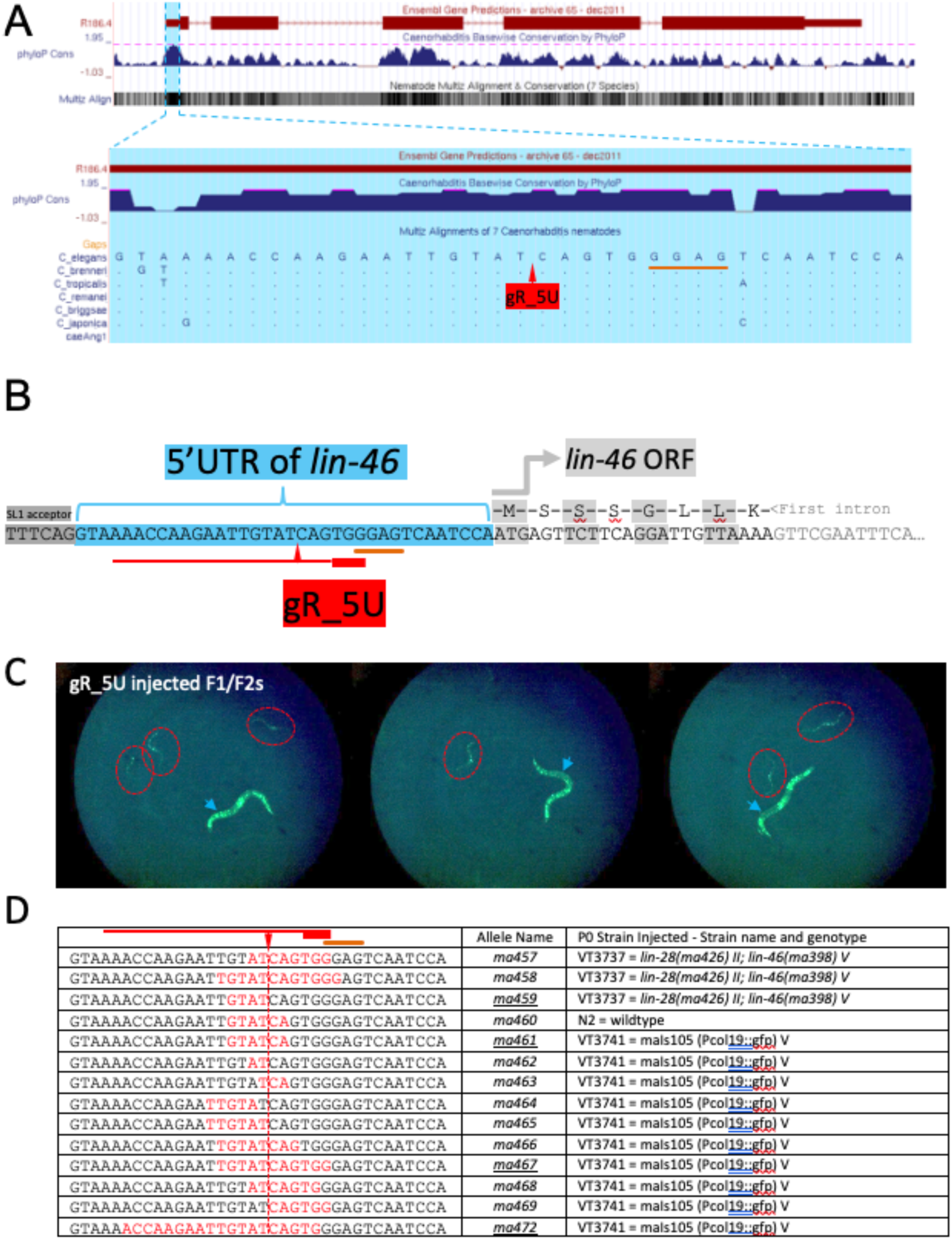
CRISPR/Cas9 mutagenesis of the conserved 5’UTR of *lin-46*. A) Genome browser view of the *C. elegans lin-46* gene (top) and the magnified 5’UTR sequence (bottom). Genome browser tracks: Ensemble Gene Predictions (top), PhyloP conservation (middle), Nematode Multiz Alignment (bottom). Note that the phyloP and Multiz tracks show the high conservation in the 5’UTR among the seven nematode species listed in the figure. The gR_5U guide cut-site and the GGAG in the 5’UTR of *lin-46* are marked. B) *lin-46* 5’UTR flanked by the TTTCAG splice acceptor and the *lin-46* ORF as well as the gR_5U and the GGAG are shown. C) Examples of F1/F2 animals among the progeny of gR_5U injected P0s that express precocious *Pcol-19::gfp* are marked with red circles. F1/F2 progeny displaying wildtype *Pcol-19::gfp* pattern are marked with blue arrowheads. D) Various mutations in the 5’UTR of *lin-46* that are detected in the F1/F2 progeny of the gR_5U injected animals displaying precocious *Pcol-19::gfp* or protruding vulva phenotypes are listed in the left column. Red fonts indicate the deleted nucleotides in each allele. Allele names given to these mutations and the P0s strains injected are given in the middle and right columns, respectively.

**Figure S2.**
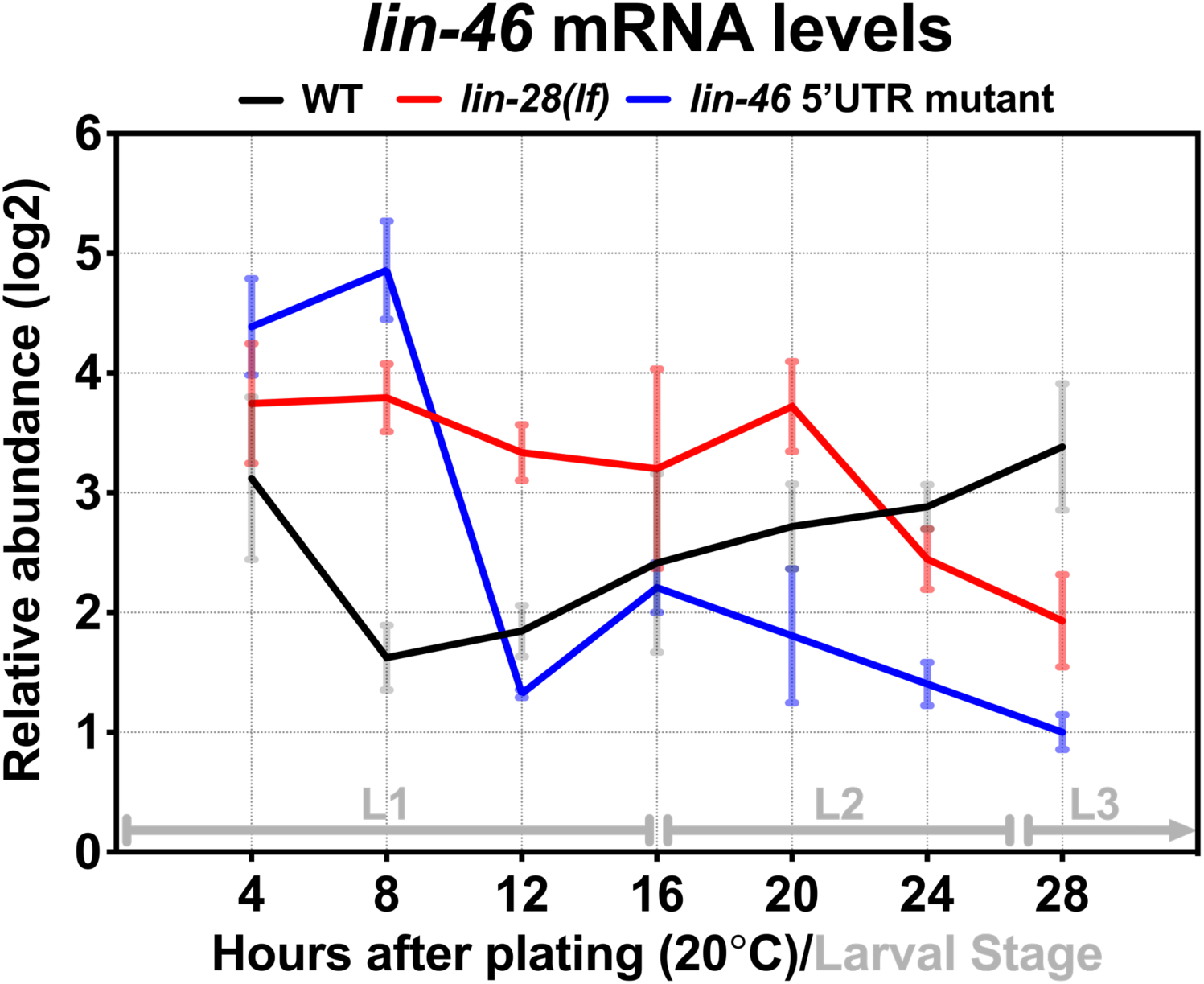
Temporal dynamics of *lin-46* mRNA expression at the L1 and L2 stages. qRT-PCR analysis of *lin-46* mRNA levels in samples of total RNA from staged populations of synchronously developing WT, *lin-28(lf)*, and *lin-46(5’UTR/ΔGTAT)* animals at 20°C. n = three biological replicates. Data are mean ± s.d.

**Figure S3.**
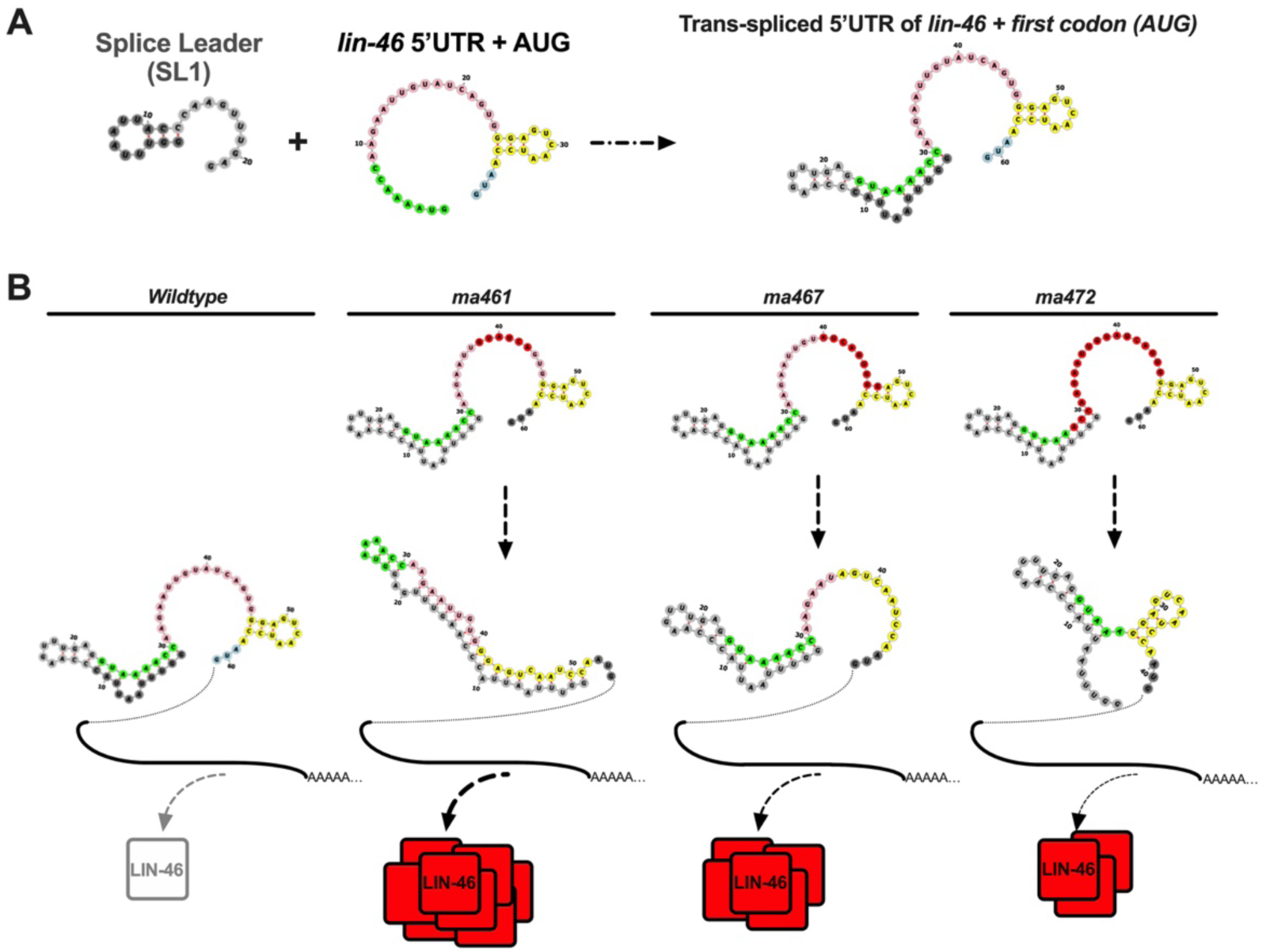
The effects of 5’UTR mutations on the folding of the trans-spliced 5’UTR of *lin-46* and corresponding LIN-46 expression levels inferred from the strength of the gain-of-function phenotypes. A) Predicted folding of the splice leader (SL1) and *lin-46* 5’UTR before and after trans-splicing. B) Predicted changes in the folding of the trans-spliced *lin-46* 5’UTR in three different mutants and schematic representation of LIN-46 expression inferred from the phenotypes (e.g. Figure 2A) observed in animals carrying each mutation.

